# Temperature-driven coordination of circadian transcriptome regulation

**DOI:** 10.1101/2023.10.27.563979

**Authors:** Bingxian Xu, Dae-Sung Hwangbo, Sumit Saurabh, Clark Rosensweig, Ravi Allada, William L. Kath, Rosemary Braun

## Abstract

The circadian rhythm is an evolutionarily-conserved molecular oscillator that enables species to anticipate rhythmic changes in their environment. At a molecular level, the core clock genes induce a circadian oscillation in thousands of genes in a tissue–specific manner, orchestrating myriad biological processes. While studies have investigated how the core clock circuit responds to environmental perturbations such as temperature, the downstream effects of such perturbations on circadian regulation remain poorly understood. By analyzing bulk-RNA sequencing of *Drosophila* fat bodies harvested from flies subjected to different environmental conditions, we demonstrate a highly condition-specific circadian transcriptome. Further employing a reference-based gene regulatory network (Reactome), we find evidence of increased gene-gene coordination at low temperatures and synchronization of rhythmic genes that are network neighbors. Our results point to the mechanisms by which the circadian clock mediates the fly’s response to seasonal changes in temperature.

**Significance Statement:** The circadian rhythm enables organisms to anticipate and adapt to changes in their environment. While behavioral changes have been observed in *Drosophila melanogaster* subjected to low temperatures, little is known regarding how these changes are enacted at a molecular level. By conducting bulk RNA sequencing from fruit flies, we observe that genome-wide circadian oscillation patterns are temperature dependent. Intriguingly, we find that morning and evening peaks of transcriptomic activity shift closer together, consistent with anticipation of a shorter photoperiod in cooler winter weather. We further find that the low-temperature dynamics are highly coordinated with respect to a reference-based gene regulatory network. Our findings provide insights into the mechanisms by which flies adapt to environmental temperature changes.

## 1 Introduction

A circadian clock is present in organisms from mold to Drosophila to mammals, where it drives the regulation of processes such as the rest/activity cycle, spore production, and metabolism [1]. In general, the circadian rhythm can be defined as a robust 24-hour self-sustaining oscillator that can be entrained by environmental cues, such as light. The molecular basis of the core clock circuit has been elucidated with great detail [1, 2, 3, 4, 5, 6]. In Drosophila, the CLOCK/CYCLE (CLK/CYC) heterodimer binds to the E-box to activate gene expression of period (*per*) and timeless (*tim*). PER and TIM proteins dimerize in the cytoplasm and translocate to the nucleus, with a time constant of approximately 4 hours, to inhibit the DNA binding activity of CLK/CYC [2]. This transcription-translation feedback loop oscillates with an approximate 24-hour period. In turn, it drives the expression levels of hundreds of downstream genes.

Recently, advances in high throughput sequencing have made it possible to identify genes under circadian control. In a study conducted by Zhang et al. [7], where they sampled 12 different mouse organs every two hours for two days using microarray, it was reported that nearly 40% of all genes showed rhythmic behavior in at least one of the twelve organs that were studied. In addition, they observed little overlap of cycling transcripts between all organ pairs, suggesting that the circadian regulation is highly tissue–specific. Nevertheless, while each organ has a largely unique set of circadian genes, their phase distribution showed much less diversity, generally peaking either in-phase or anti-phase with respect to CT20.

A hallmark of the circadian clock is the insensitivity of its endogenous period to temperature, a phenomenon known as “temperature compensation” [6]. This property is in part attributable to gene-specific thermosensitive alternative splicing [3], which preserves the period of the feedback loop even as reaction rates change. However, while the *period* remains constant across a wide range of temperatures, the *phase* of the clock has been observed to be temperature sensitive. For example, behavioral studies using Drosophila Activity Monitoring (DAM) systems have demonstrated that under normal light–dark cycles, flies show gradual increase in activity just before the light–dark/dark– light transitions, a phenomenon known as evening/morning anticipation [2]. When the temperature drops, the activity peaks show decreased separation, with an earlier onset of evening anticipation that may reflect an adaptation for the shortened daylight hours in winter.

While this temperature-dependent behavioral pattern has been observed in Drosophila, comparatively little is known about how downstream transcriptome-wide circadian activity changes in face of temperature perturbations. Early work reported that the transcriptome was modified globally by a cyclic temperature perturbation, which acted as a driver of circadian oscillation [8]. More recently, it was reported that flies subjected to different environmental temperatures under the light–dark cycle exhibited thermosensitive alternative splicing, affecting locomotor activities [9]. Taken together, these works provide evidence that temperature could have a global effect on the circadian transcriptome. However, both aforementioned studies had a sampling frequency of four hours, which hinders the detection of cycling genes and the estimation of their associated phases [10].

To further understand how temperature impacts the circadian transcriptome, we conducted genome-wide studies of circadian gene expression in fruit flies subjected to a 25*^◦^*C or 18*^◦^*C environment by performing bulk RNA sequencing on fat body samples with a sampling frequency of two hours. The fat body in Drosophila is a major endocrine organ responsible for metabolism and energy storage [11]. It has been observed that genes cycle with a 24-hour period in the fat body, with consequences for metabolism [12]. However, it is not known whether prolonged temperature change can induce phase shift, or initiates circadian oscillation of “flat” genes. By conducting bulk RNA-sequencing, we show that both the identity and phases of cycling genes change in a temperature–dependent manner. In addition, we report a significant increase at lower temperatures in the phase synchronization of oscillating genes that are close to one another on the Reactome Drosophila gene regulatory network, suggesting that transcriptomic coordination may be enhanced at lower temperatures.

## 2 Results

### Circadian transcriptional profiling

To study how the circadian rhythm is perturbed by temperature, we conducted two separate studies, henceforth termed V1 and V2. In experiment V1, Drosophila fat bodies were harvested every two hours in 12:12 L:D conditions after flies had been at 25*^◦^*C and 18*^◦^* for four days (see Figure 1), providing data for a stable, established circadian transcriptome. In the V2 experiment, in addition to examining the stable circadian transcriptome during days four and five, we also collected data during the first two days, investigating its transient dynamics. Analysis of the transient data is reported separately [13]; here, we focus on the dynamics after flies had equilibrated to the 18^◦^C and 25^◦^C conditions. These two separate experiments provide the opportunity to minimize the chances of falsely identifying cycling genes and ensured our results are robust to sequencing technology and experimental artifacts, which can be a major source of error in cycling detection [10, 14].

**Figure 1:**
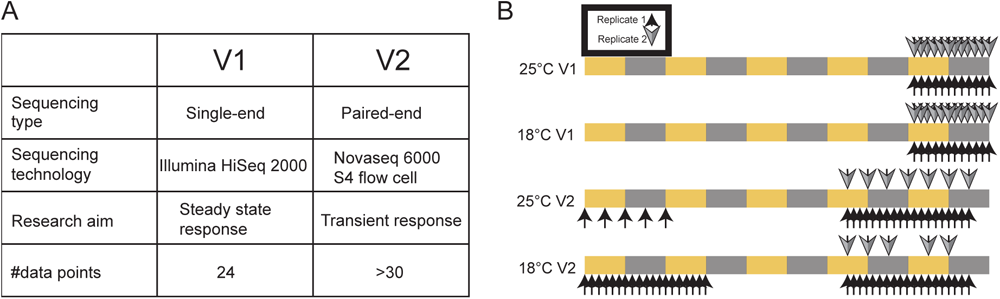
Experimental design. A: Method and technologies used for the V1 and V2 experiments. B: Sampling scheme of the two experiments and environmental conditions. Arrows above and below the colored bars indicate samples and replicates. The colored bars depict the light (yellow) and dark (grey) periods.

**Figure 2:**
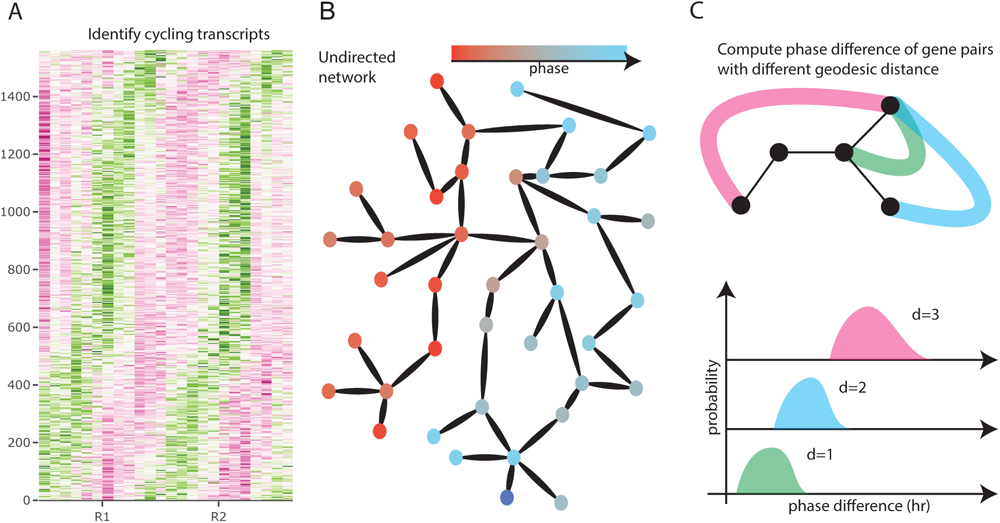
Illustration of the analysis pipeline. Circadian genes are identified under each temperature (A). Estimated phases phases were mapped onto a database–derived network (B). From the network, we look for evidence of phase organization by quantifying phase difference as a function of geodesic distance on the network (C).

To understand how the circadian transcriptome is affected by temperature, we first identified cycling genes at 25*^◦^*C and 18*^◦^* and estimated their phases (Figure 7.2A). Next, we investigated whether there is evidence of increased synchronization of gene expression dynamics at different temperatures, by examining phase differences between genes in the context of putative gene regulatory networks. We obtained the gene regulatory network from the Reactome database [15] (Figure 7.2B), an expert–curated database of experimentally–validated Drosophila–specific gene–gene interactions, and tested whether genes with similar phases were clustered on the graph (Figure 7.2C).

### Identification of rhythmic transcripts

It has been shown that cycling detection is a challenging task, depending upon sampling resolution, cycling detection algorithm, and the dataset. To deal with these problems, we employed two strategies. First, we used harmonic regression to test for evidence of cycling, assigning the *F* –test *p*-value to each gene. We did this for both the V1 and V2 experiments, and considered a gene to be cycling only if (i) it has a harmonic regression *p <* 0.1 in *both* experiments and (ii) the estimated phase difference between the two experiments Δ*ϕ <* 3*h*, indicating consistency between the two experiments. We also compared the estimated phases and *p*-values assigned by harmonic regression to those obtained from JTK-CYCLE [16] and observed a high concordance (Supplementary Figure S1A, B). Prior to the cycling detection step, we filtered out genes that have low expression or are rarely detected (see Materials and Methods for detail), retaining 6774 genes. With these criteria, we would expect 6774 *×* 0.1^2^ *×* <u>^3^</u> = 17 genes to be cycling be chance. In our data, we identified 242 and 364 cycling genes in 18*^◦^*C and 25*^◦^*C respectively, of which 79 overlapped (Figure3A, Supplementary Figure S2, Supplementary table 1). Of these, only 17 — 7% and 4.7% respectively for the two temperatures — are expected to be be false discoveries (Supplementary Figure S3A, B).

As an additional validation, we examined the dynamics of the core clock genes (*tim*, *Clk*, *vri*, *pdp*, *per*, *cry*) and observed that their estimated phases were highly concordant between the V1 and V2 experiments (Supplementary Figure S4A). In addition, we noted a phase advance of the core clock genes at 18*^◦^*C relative to 25*^◦^*C (Supplementary Figure S4B).

Taken together, this analysis suggests that, first, oscillations are induced in a condition specific manner; and second, a phase advance occurs in the core clock genes at the lower temperature, potentially influencing its downstream targets.

### Functional analysis of cycling genes

To examine the functionality of cycling genes in each condition, we conducted an overrepresentation analysis on genes that cycle in 18*^◦^*C, 25*^◦^*C, and at both temperatures using the ReactomePA package [17] in R. The significantly over-represented pathways in 18*^◦^*C and 25*^◦^*C showed little overlap (Supplementary tables 2,3). At 18*^◦^*C we found that the most over-represented circadian pathway is the metabolism of lipids (*q* = 0.003). In addition, many lipid-related metabolic pathways are significantly over-represented as well, such as triglyceride metabolism (*q* = 0.006), fatty acid metabolism (*q*=0.006), carnitine metabolism (*q* = 0.02). At 25*^◦^* however, we observed only three significantly over-represented pathways: pentose phosphate pathway (*q* = 0.006), metabolism of amino acids (*q* = 0.04) and derivatives, and ABC family proteins mediated transport (*q* = 0.04). For genes that were cycling at both temperatures, we observed that the circadian clock pathway is over-represented as expected (*q* = 0.017; Supplementary Table 1). In addition, we found that metabolism of lipids to be an over-represented pathway (*q* = 0.018).

Through deeper investigation, we found that there were 28 and 27 cycling genes involved in the metabolism of lipids at 25*^◦^*C and 18*^◦^*C respectively, with only 13 genes cycling under both temperatures. In addition, for the condition–specific cycling genes within the lipid metabolism pathway, we found that their average *p*-values in the other condition were close to 0.4; that is, the subset of genes driving the significance of the lipid metabolism pathway are highly temperature–specific, with significant evidence of cycling in one condition and far from significant in the other. Together, these results suggest that temperature affects the lipid metabolism in the fat body, and that there are mechanisms that exercise precise control of rhythmicity even within the same pathways.

### Distribution of Phases

To characterize cycling genes further, we estimated their phases at 25*^◦^*C and 18*^◦^*C individually by taking the circular mean of phases estimated from the V1 and V2 experiments. Interestingly, we found that the phase distributions of all detected cycling genes at both temperatures are significantly different (*p* = 0.04, KS test) and were both bimodally distributed, peaking at ZT 4.6 and ZT 20.6 at 25*^◦^*C and at ZT 7.6 and ZT 20.6 at 18*^◦^*C (Figure 3B). The narrower interval between gene expression activity peaks at the lower temperature is analogous to the previous observation that activity peaks are closer to each other at low temperature [3].

**Figure 3:**
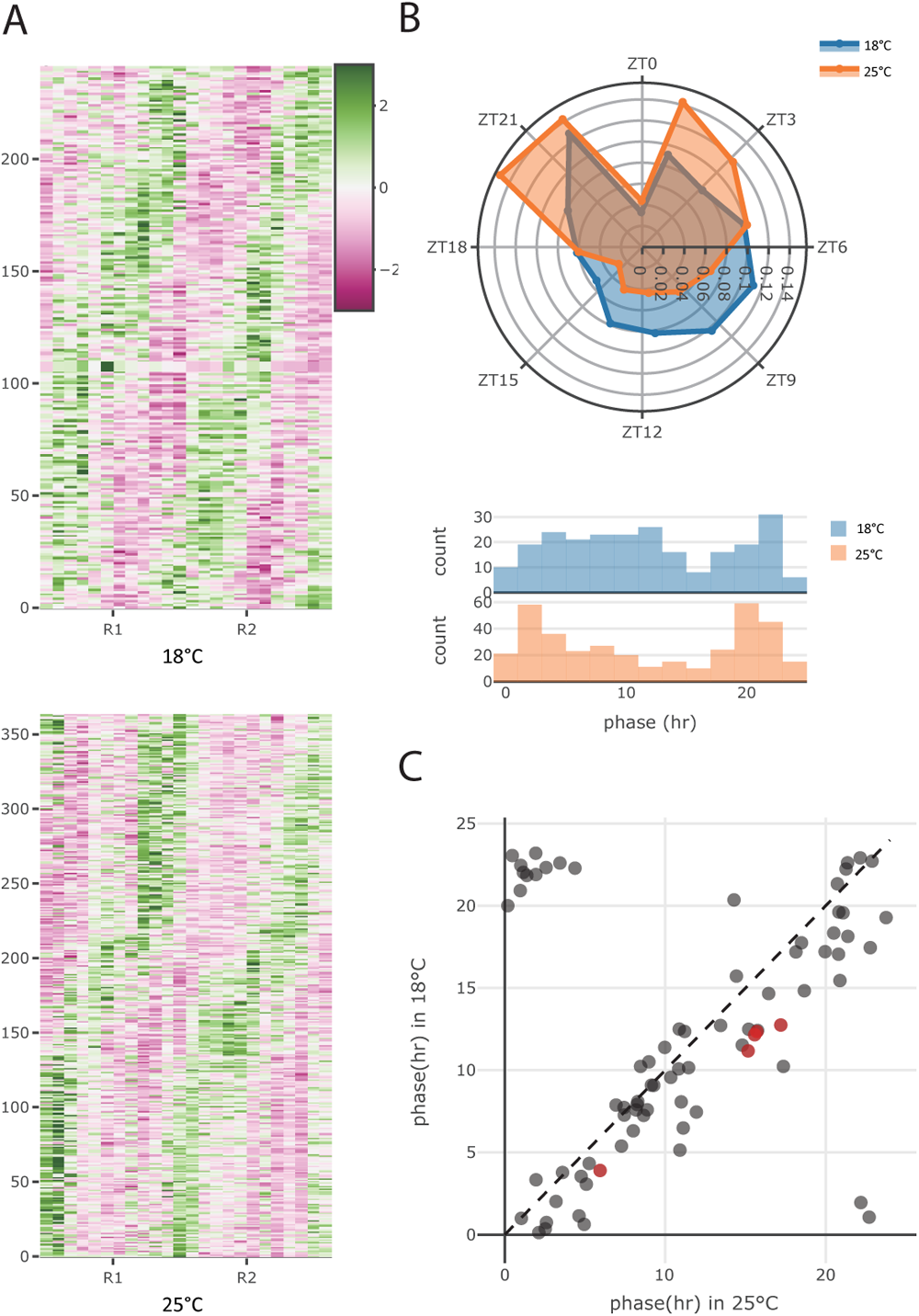
Phase distributions. A: *Z*-scored TPM of detected cycling genes in the V1 experiment. (For the V2 experiment, see Supplementary Figure S2.) B: Roseplot (top) and histogram (bottom) of the phase distribution for all genes cycling at either 18*^◦^*C (blue) or 25*^◦^*C (orange). C: Estimated phases of genes that cycle at both temperatures. Red dots highlight the core clock genes.

To examine how rhythmic expression was perturbed by temperature, we investigated the genes that were considered cycling at both temperatures (n=79) (i.e., the intersection of all cycling genes) and found their phases to be highly similar, having a circular correlation [18] of 0.79 (Figure 3C). Interestingly, for genes that were considered cycling in both temperatures, we observed an almost universal phase advance in 18*^◦^*C relative to 25*^◦^*C (Figure3C) and found that the extent of this phase advance is concordant in V1 and V2 experiments using limorhyde [19, 20](Supplementary figure S5), having a mean phase shift of 2.02 and 1.36 hours respectively. Interestingly, the phase shift for the common cyclers appears to be the same regardless of the phase of the gene; that is, among the common cyclers, we do not observe the only the morning–peaking genes shifting phase. This implies that the differential change in the peaks of gene activity observed in Figure 3B are attributable to condition–specific cyclers.

From our results we speculated that environmental perturbation can impact circadian transcriptome in two ways. First, oscillation can be induced in a condition-specific manner, as has been observed in flies subjected to our two experimental conditions. Second, temperature perturbation impacts another set of rhythmic genes, initiating a universal phase advance, potentially mediated by the core clock genes.

### Network Phase Organization

We next examined phase organization with respect to the gene regulatory network obtained from the Reactome database [15] via the R Graphite library [21, 22]). Since the network only contained a subset of the genes, we first checked whether these were representative of all genes by testing whether the phase distribution of genes on the network differed from that of all detected cycling genes. We found that the Reactome genes did not have a significantly different phase distribution relative to all genes (Kolmogorov-Smirnov test, 18*^◦^*C: *p* = 0.9778, 25*^◦^*C: *p* = 0.6806), suggesting that these are a representative sample. We then tested whether the connectivity of a gene is predictive of its rhythmicity. Comparing the degree distributions of 18*^◦^*C and 25*^◦^*C cycling genes to that of all genes, we observed that these distributions were not significantly different from one another (Supplementary Figure S6A).

We then examined whether cycling gene are co-localized on the network by computing pair-wise distances between 18*^◦^*C and 25*^◦^*C cycling genes separately, compared these distance distributions to that obtained from all genes. We found that the identified cycling genes at both 18*^◦^*C and 25*^◦^*C tend to have smaller network distances than expected from all genes on the network (18*^◦^*C cyclers vs all: *p* = 2 *×* 10*^−^*^16^; 25*^◦^*C cyclers vs all: *p* = 2 *×* 10*^−^*^16^, Wilcoxon rank sum test, Supplementary figure S6B), suggesting localization of cycling genes with respect to the graph.

If it is the case that cycling genes are coordinated by the gene regulatory network, we may expect that genes that are closer together on the network will exhibit smaller phase differences than those that are farther apart. To test this idea, we quantified how the association between phase differences between gene pairs and their geodesic distances on the network (Figure 4A). We found that the phase difference distribution computed at the two temperatures exhibits a distinct pattern. While the median phase difference remained approximately constant at 25*^◦^*C (slope = 0.025, *p* = 0.69, linear regression), it steadily increased with geodesic distance at 18*^◦^*C (slope = 0.30, *p* = 0.019, linear regression). To further characterize this pattern, we constructed a null models by randomly reassigning the locations of cycling genes on the network 5000 times (Figure 4B). Compared to the null model, the phase differences of immediate neighbours at 18*^◦^*C were significantly smaller than that of any pair of genes selected by chance (*p* = 0.018, Figure 4B). At 25*^◦^*C however, none of the median phase differences were significant.

**Figure 4:**
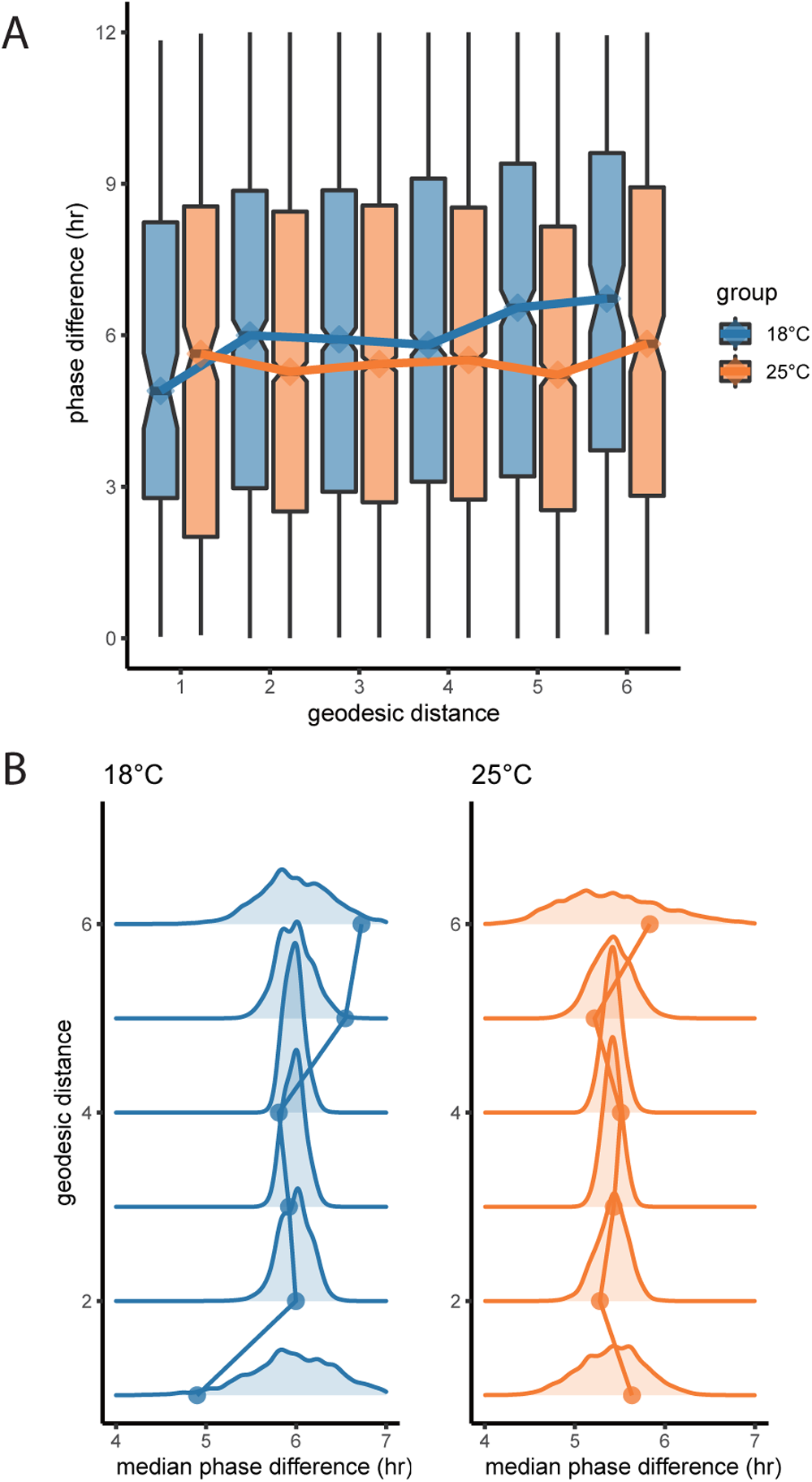
Temperature perturbation alters circadian gene expression globally. A: Distribution of phase differences of gene pairs with at given geodesic distances. Lines connect the median under the two temperatures. B: Observed median phase difference (points) at each geodesic distance compared to the null distribution (curves) of expected phase differences at each distance.

We considered the association between network position and phase to be evidence of phase organization across the network. We observed this in the low temperature condition only, and we reason that there are several plausible explanations: either we have reduced ability to detect cycling genes and estimate phases at 25*^◦^*C due to increased transcriptional noise or increased cell-cell desyncrony, or there is a true biological reprogramming of the circadian transcriptome. To investigate these possibilities, we conducted two additional analyses.

First, increase in temperature could increase transcriptional noise, hindering our ability to detect cycling genes at a higher temperature. To test this, we exploited the fact that most genes do not show rhythmic behavior, quantified the gene expression variance for all genes in all four datasets (two from V1 and two from V2), shown in Supplementary figure S7A. We found that gene expression variance was higher at 25*^◦^*C than 18*^◦^*C in the V1 experiment (*p* = 0.0005, two tailed Wilcoxon signed rank test), but (strongly) higher at 18*^◦^*C relative to 25*^◦^*C in the V2 experiment (*p* = 2*×*10*^−^*^16^, two tailed Wilcoxon signed rank test). When the variances from V1 and V2 are combined, the increased variance at 18*^◦^*C relative to 25*^◦^*C in V2 dominates, such that 18*^◦^*C appears to have higher variance overall (*p* = 2 *×* 10*^−^*^16^). This result, and the lack of consistent association between V1 and V2 suggests that there is no evidence for increased variance at higher temperatures.

Alternatively, we might also observe a loss of rhythmicity if cells become asynchronous under higher temperature, effectively damping the oscillations that could be observed in the bulk; because our data were collected using bulk RNA sequencing, successful detection of rhythmic activity requires most cells to oscillate synchronously. A loss of synchrony amongst cells would result in a lower amplitude observed in the bulk, even amongst cycling genes. We thus tested whether the amplitude of genes that were detected as cycling under both temperatures exhibited reduced amplitude at higher temperatures (Supplementary figure S7B). We observed an decreased oscillation amplitude at 25*^◦^*C relative to 18*^◦^*C in the V1 experiment (*p* = 5 *×* 10*^−^*^5^, two tailed Wilcoxon signed rank test), but an increased oscillation amplitude at 25*^◦^*C relative to 18*^◦^*C in the V2 experiment (*p* = 0.017, two tailed Wilconxon signed rank test). Combining the V1 and V2 amplitudes negates any association between temperature and amplitude (*p* = 0.2). As before, we observed no consistent association between oscillation amplitude and temperature, implying that the tissue is likely equally well synchronized at 18*^◦^*C and 25*^◦^*C.

These results suggest that the effect we observe (increased cycling and phase organization at 18*^◦^*C) is due to a reprogramming of the circadian transcriptome. By considering oscillation amplitude and gene expression variance, we reason that the lack of phase organization at 25*^◦^*C is not a consequence of either increased transcriptional noise or loss of cell-cell synchronization. The results suggest that phase coordination across the gene regulatory network is important for survival under harsher environmental conditions, and this is manifest as circadian phase organization at 18*^◦^*C.

## 3 Discussion

By conducting bulk RNA sequencing at different temperatures, we have found that genes in *Drosophila melanogaster* are induced to cycle in a temperature–dependent manner. Examining the localization of cycling genes on a gene regulatory network, we detected statistical evidence of assortativity (clustering of similar phases on the network), and thus hypothesize that rhythmic gene expression is organized along the network. To test this idea, we quantified the dependence of phase difference of gene pairs on geodesic distance, confirming the presence of phase organization on the network in the low temperature condition. We also found that the differences in cycling primarily affected the activity of pathways associated with lipid metabolism, suggesting that the fruit-fly uses the circadian clock to mediate its adaptation to adverse (i.e. lower temperature) conditions. The fact that the morning peak of gene expression activity shifted later at lower temperatures, consistent with a shorter winter day, may be evidence for a program of seasonal adaptation in the fly. That is, at lower temperatures, the fly not only changes its metabolic activity, it also changes *when* that metabolic activity happens. We emphasize here that the flies continued to receive 12:12 L:D, and thus this change was driven by temperature alone.

Existing studies of the circadian oscillations have focused primarily on the identification of rhythmic transcripts [7] or the functionality and impact of a small group of genes under circadian control [23, 24]. However, these studies have have not shed light on the potential impact of circadian oscillations due to gene-gene interactions. By adopting a network-based approach, which allows direct observation of collective behavior of circadian genes in conjunction with environmental or genetic perturbation, one observes this additional aspect of circadian regulation.

We acknowledge that our study only considers one type of environmental perturbation, namely a low-temperature perturbation. To fully understand how circadian control changes with environment, it would be important to conduct similar studies under a wider range of conditions. For example, it would be valuable to conduct similar experiments at a range of temperatures in order to investigate how cycling and phase organization changes under finer temperature changes. It may also be of interest to investigate other tissues, as circadian synchronization and entrainment is known to differ between central and peripheral clocks [5, 25, 26]. Additionally, our study only probed transcriptomic changes; temperature–dependent post–translational modifications could not be probed here but would give deeper insights into the effect of temperature on metabolism.

Finally, an open question is how quickly the genes change their cycling behavior to adapt when the temperature changes. Observing the transient dynamics of gene expression activity immediately following the temperature change can yield insights into this question. For investigations of this question using the V2 data, the reader is referred to [13].

## 4 Materials and Methods

### Fly Rearing and Media

All the flies used for experiments were raised on a standard yeast-cornmeal-molasses based diet under a light-dark (LD) 12:12h cycle. All experiments were carried out with Iso31 flies (Bloomington Drosophila Stock Center ID: 5905). Age matched female flies were collected and aged for 3 days with males before being transferred to an entrainment incubator.

### Fat Body Dissection

Young mated female flies (*∼*3 day old) were entrained under a control diet (15SY) for 5 days in 12L:12D cycles at 25*^◦^*C. Every 2 hours, flies were directly dissected without dry ice to harvest fat tissues in the abdomen. Pinned flies were cut to remove organs in the abdomen (intestine, ovaries, malpighian tubules, etc.). Fat tissue attached to epidermis was collected (1-3). Fat body from *∼*10 flies were harvested within 10 minutes for each time point of RNA-Seq analysis.

### 4.1 RNA extraction

RNA was isolated from the abdominal fat bodies using Trizol LS (ThermoFisher, #10296028). Briefly, 300 *µ*L of Trizol LS was added to fat bodies in 100 µL of PBS. The tissue was homogenized with a motorized pestle for 2 minutes before adding another 600 *µ*L of Trizol LS (3:1 mixture of Trizol LS:PBS). The resulting solution was centrifuged at 12,000 x g for 10 minutes at 4*^◦^*C. The aqueous supernatant layer was collected in a new tube, while carefully avoiding disturbing the other layers of the phase separated solution. RNA was extracted from the aqueous supernatant layer by vigorously shaking with 240 µL of chloroform (Fisher Scientific, #C298), again carefully avoiding other layers following phase separation. The aqueous phase was transferred to a new tube and the RNA was precipitated by incubating at room temperature with 500*µ*L of 100% isopropanol (Sigma-Aldrich, #I9516). Following centrifugation at 12,000g for 10 minutes at 4*^◦^*C, the supernatant was removed leaving only the RNA pellet. The pellet was washed with 1 mL of 75% ethanol (Sigma-Aldrich, #E7023), then air dried for 5-10 minutes before resuspension in RNase-free water.

### RNA-seq V1

cDNA library was constructed with poly(A) selected mRNA using Truseq RNA library preparation kit and then sequenced at the Genomics Core Facility at the University of Chicago on Illumina HiSeq 2000 System.

### RNA seq V2

Purified RNA was sent to Novogene (Sacramento, CA) for library preparation. Libraries were prepared from mRNA purified from total RNA using poly-T oligo-attached magnetic beads (NEBNext Ultra II RNA Library Prep kit for Illumina, New England Biolabs, E7775). Non-stranded library preparation was carried out using the NEBNext Ultra II RNA Library Prep kit for Illumina according to manufacturer protocol. Libraries were subsequently sequenced on a Novaseq 6000 S4 flow cell. 20 million paired-end reads (PE150) were generated for each sample.

### Read alignment

Bulk RNA seq data were first quality checked using FastQC [27] and trimmed using Atropos [28]. Reads (deduplicated for V2) were aligned and quantified using STAR [29] and RSEM [30] to the Ensembl *Drosophila melanogaster* BDGP6.32 reference (release 107) using standard parameters (See supplementary file for detailed protocols).

### Data preprocessing

We selected genes with median TPM*>* 5 in either the 18*^◦^*C or 25*^◦^*C condition, consistently across the two experiments. 6774 genes passed this filter in total, and used these genes only for further analysis.

### Quantification of phase difference

The phase difference Δ*ϕ/* between two circular variables is defined as min(*θ*_1_*, θ*_2_) where:

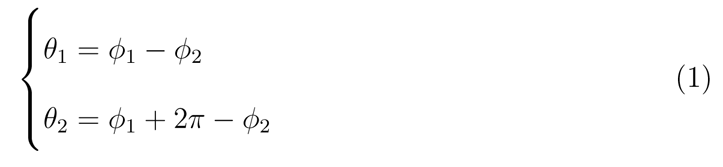

Here, *ϕ*_1_ and *ϕ*_2_ are the angular phase, in radians, of detected cycling genes and we assign them such that *ϕ*_1_ *> ϕ*_2_.

### Identification of rhythmic genes

Harmonic regression was used to identify cycling genes and estimate phases. For each gene, we have two *p* values and phase estimates, one from the V1 and one from the V2 experiment. A gene is considered to be cycling if both the *p <* 0.1 in *both* V1 and V2, and the phase estimates differ by no more than three hours (Δ*ϕ <* 3*h*).

Although we used harmonic regression to conduct cycling detection, which can be vulnerable to false positives, we reasoned that our combination of the V1 and V2 data mitigated this concern. Furthermore, we compared the performance of harmonic regression and JTK-CYCLE and observed that the two methods produce the same phases and correlated *p* values.

### Differential cycling analysis

Differential cycling analyses were conducted using the limorhyde [19] and the limorhyde2 [20] package in R following its standard procedures with default parameters.

### Network analysis

The network used in our analysis was constructed using the graphite package in R [21, 22]. We first constructed a graph of all genes using the known information from the Reactome [15] database given in the graphite package. To facilitate later analysis, the largest connected component within this graph of all genes was extracted. Given our interest in understanding phase organization, small disconnected networks will not be as informative, hence this largest component was used for subsequent analysis. We note, furthermore, that the largest connected component comprises the majority of genes and edges in the graph. The original network contained 4080 nodes and 212,808 edges; the largest connected component contained 3875 nodes and 205,758 edges.

### Over-representation analysis

Over-representation analysis of Reactome genes was conducted using “enrichPathway” function in clusterProfiler package in R [31, 17] with default parameters. For GO analysis, we used the “enrichGO” function in the same package with the same default parameters. All genes that passed the median threshold were used as the universe input.

## 5 Acknowledgements

This work was supported by NSF grant DMS-1764421 and Simons Foundation grant 597491.

This work was also supported by Defense Advanced Research Projects 520 Agency (DARPA) (D12AP00023 to RA). This effort was in part sponsored by DARPA; the 522 content of the information does not necessarily reflect the position or the policy of the government, 523 and no official endorsement should be inferred.

DSH was supported by the 528 National Institutes of Health T32 Institutional Training Grant (Northwestern Univ: grant NIH 529 T32HL007909).

## 6 Data and Code Availability

The transient data with temperature step change is accessible at NCBI GEO accession number GSE241002. The other steady-state data used for gene selection is accessible at NCBI GEO accession number GSE241003. Code for our analysis is available on bitbucket (https://bitbucket.org/biocomplexity/phase_org_figures)

## 8 Supplemental material

### Comparison against JTK-CYCLE

JTK-CYCLE was used to validate the performance of harmonic regression. For the V2 experiment, replicates were averaged first before the standard JTK-CYCLE protocol with period 24 hr and sampling interval 2 hrs. *p* values computed from JTK-CYCLE were highly correlated with those computed using harmonic regression (log linear regression: V1 25*^◦^*C slope: 0.81; V1 18*^◦^*C slope: 0.85; V2 25*^◦^*C slope: 0.69; V2 18*^◦^*C slope: 0.80; supplemental Figure S1B).

### Read alignment

Basic quality checking of sequence files was performed with FastQC [27]. Paired-end reads were first trimmed using Atropos version 1.1.31 [28] using the options

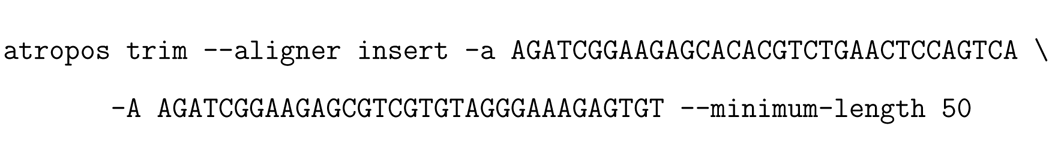

Reads were then aligned and quantified using STAR [29] (version STAR 2.7.10a alpha 220818) and RSEM [30] (version 1.3.1). STAR and RSEM indexes were first built using the Ensembl *Drosophila melanogaster* BDGP6.32 reference (release 107) using standard parameters. STAR was used with the options

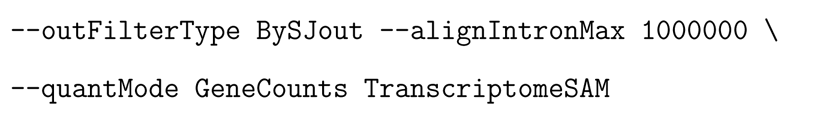

to produce raw counts and also a BAM file with reads aligned to transcriptome. RSEM was then used with options --paired-end --strandedness none to produce tags-per-million (TPM) counts for each gene from transcriptome alignments. Postprocessing of the count data into table form was performed with custom Perl, Python and shell scripts.

Because FastQC reported significant sequence duplication in the samples, we also performed the same analysis as above after deduplicating the reads. First, a BAM file was created using STAR with the options

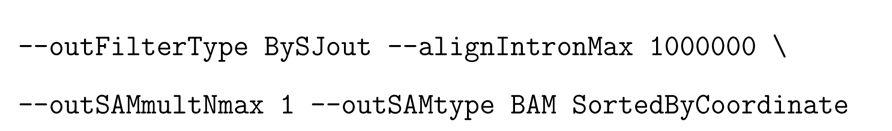

to produce only uniquely mapped reads, and then duplicate paired-end reads were removed using bamUtil [32] with the options

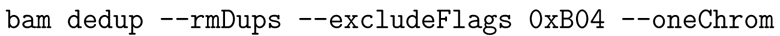

The resulting reads were then re-aligned with STAR to produce a BAM file with reads aligned to the transcriptome, followed by RSEM with the same options as above to produce TPM counts. De-duplication significantly decreased the number of reads with large TPM values (e.g., *>* 100), and as the result the TPM values of genes with smaller TPM values were increased by approximately a factor of 1.6-1.7. Deduplicated reads were used throughout the paper.

Single-end reads were quality-assessed with FastQC and then trimmed with Atropos using the options

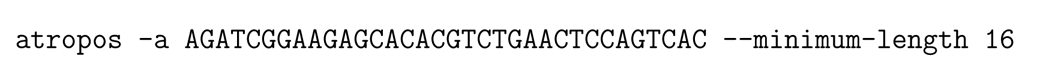

Reads were then aligned and quantified using STAR with the same options as before,

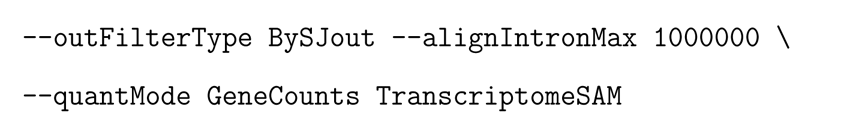

followed by RSEM with the options --strandedness none.

**Figure S1:**
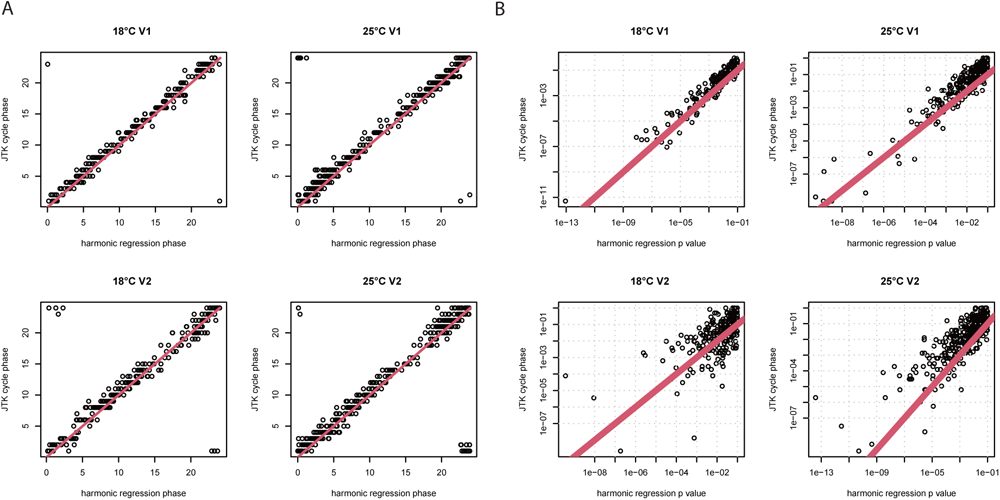
Comparison of harmonic regression and JTK-cycle. A: Phases of cycling genes estimated using harmonic regression and JTK-CYCLE. B: *p* values estimated using harmonic regression and JTK-CYCLE.

**Figure S2:**
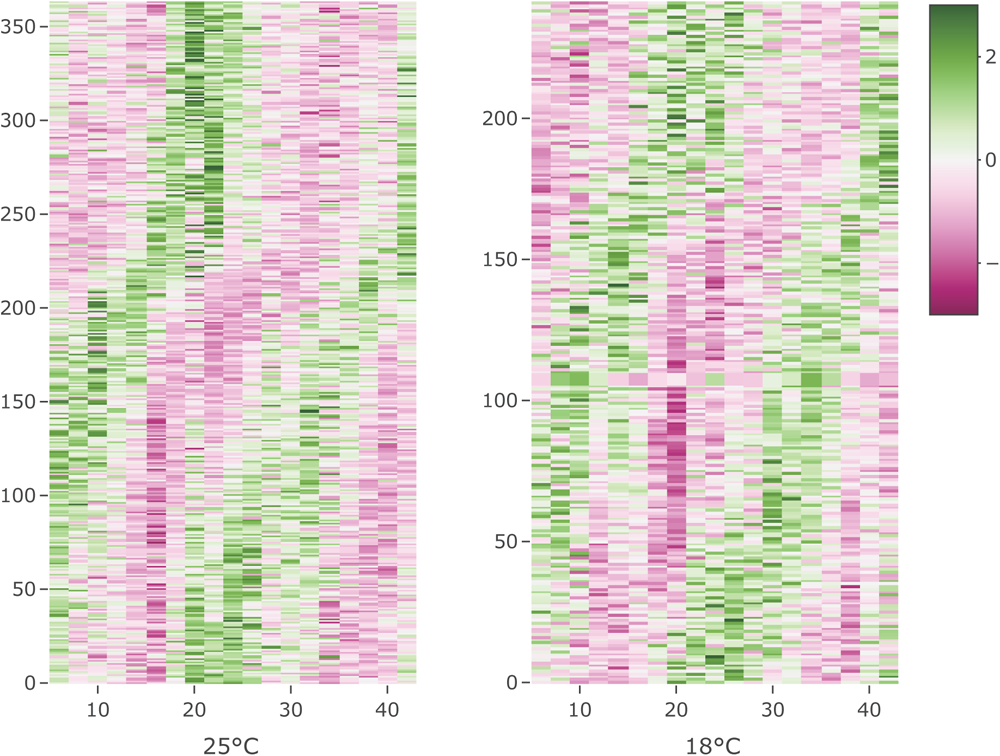
Heatmap showing the *Z*-scored TPM of the identified cycling genes in the V2 experiment. For visualization purposes, replicates were averaged.

**Figure S3:**
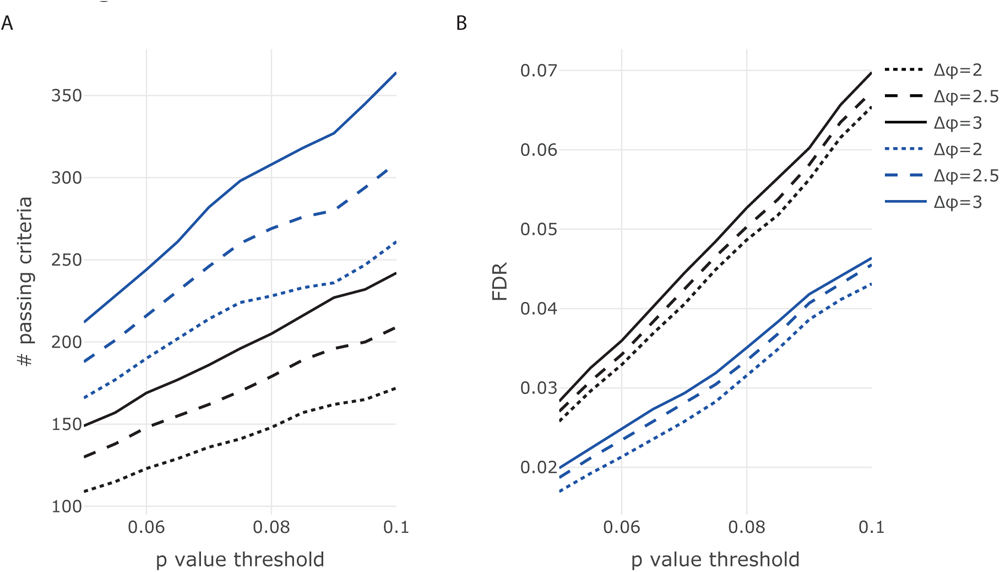
Number of identified cycling genes (A) and false discovery rate (B) as a function of harmonic regression *p*-value thresholds under different Δ*ϕ* thresholds. Blue: 25*^◦^*C. Black: 18*^◦^*C. Our selected thresholds, *p <* 0.1 and Δ*ϕ <* 3 yields FDRs of 0.07 and 0.047 in 18*^◦^*C and 25*^◦^*C, respectively.

**Figure S4:**
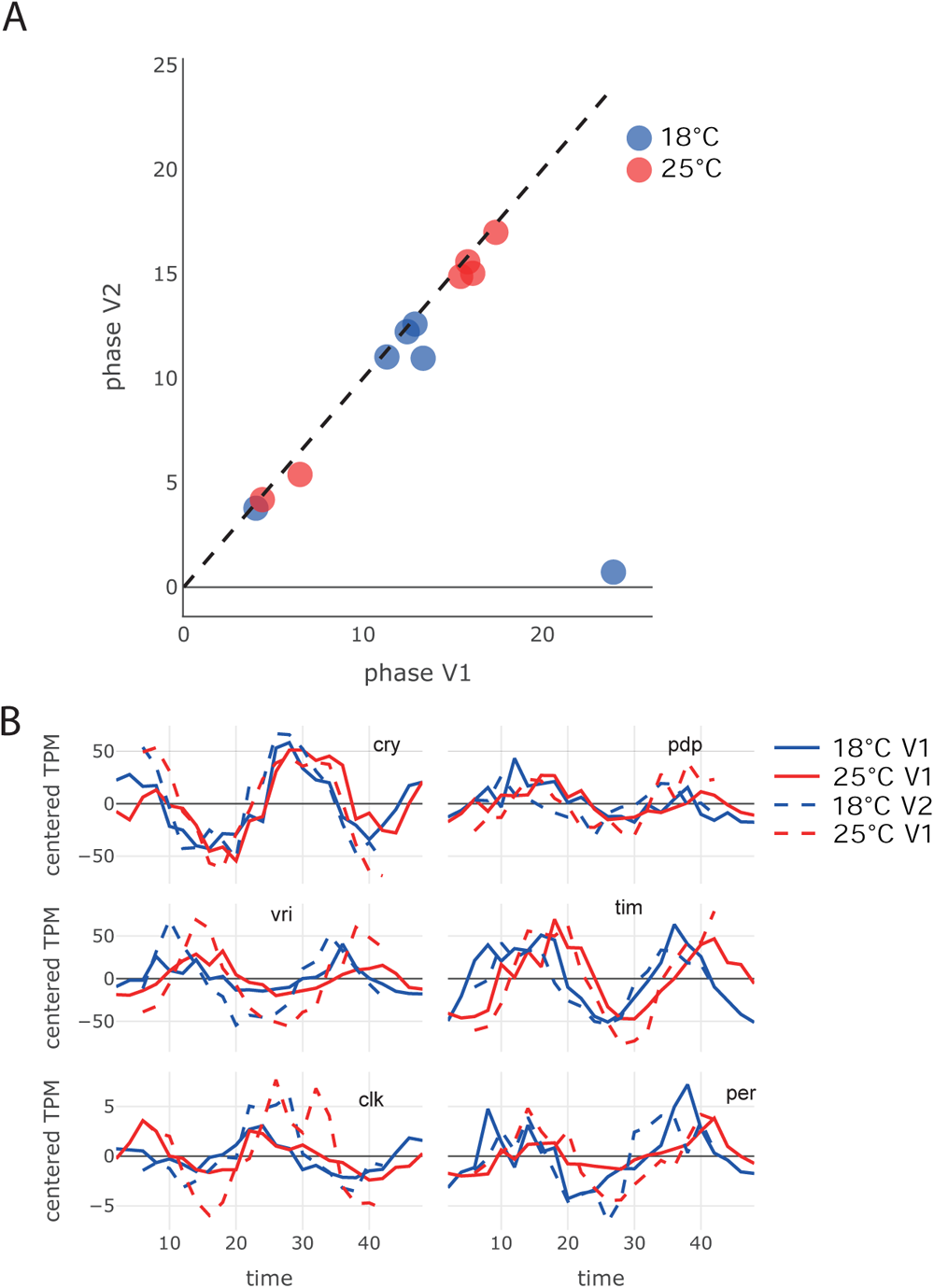
A: Phases of the core clock genes estimated in the V1 and V2 experiments, dashed line indicates *y* = *x*. C: Centered TPM of core clock genes. Replicates were concatenated for V1 and averaged for V2 for better visualization.

**Figure S5:**
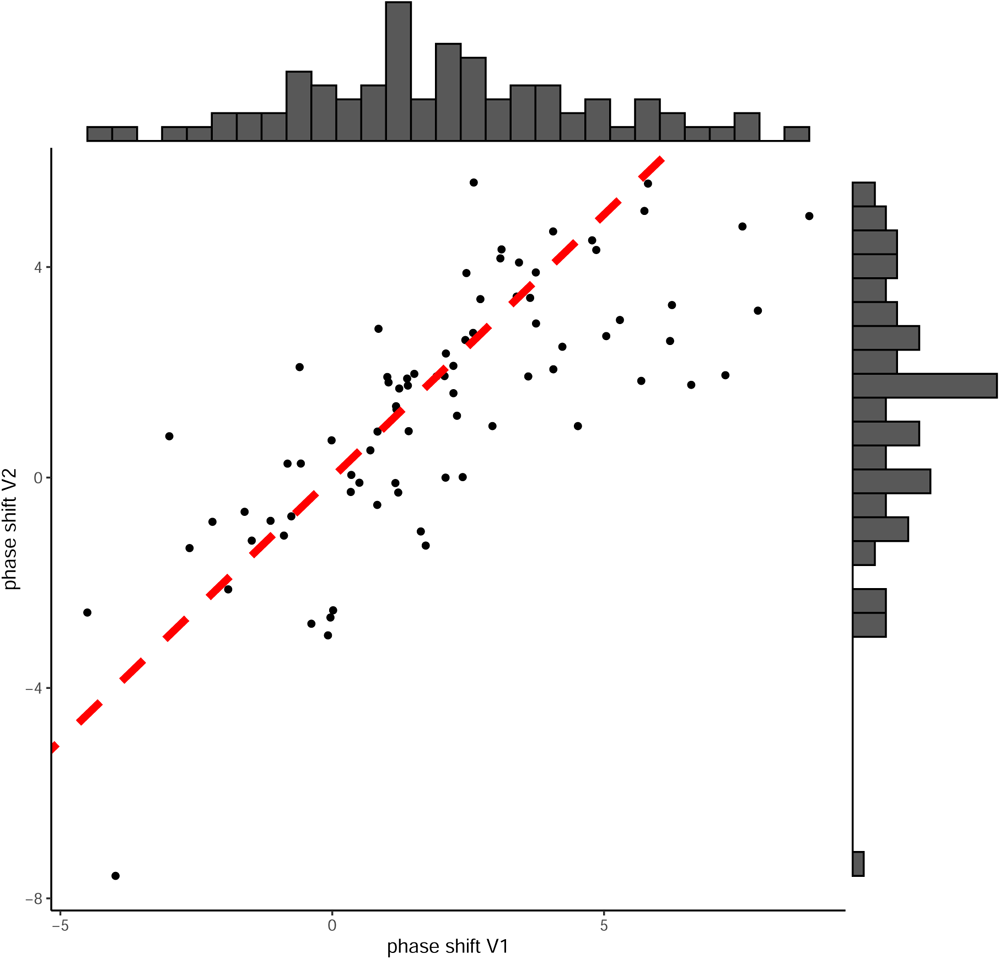
Estimated phase shift of genes that cycle under both temperatures (estimated via limorhyde2). Red line indicates *y* = *x*

**Figure S6:**
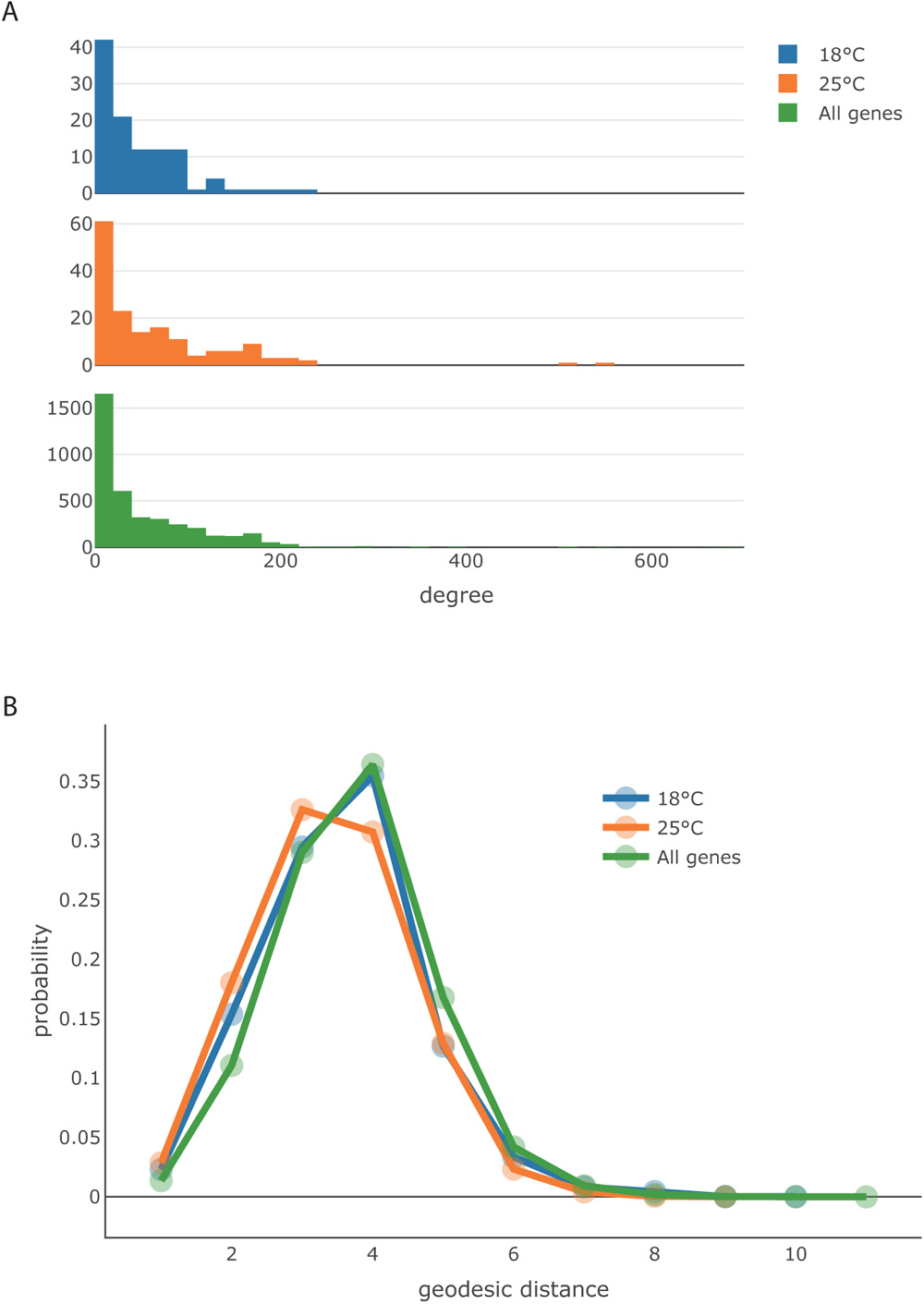
Network properties of cycling genes. A: Degree distributions of genes detected as cycling under the two temperatures, as well as all genes. B: Geodesic (network) distance distributions for gene pairs that are cycling under the two temperatures, as well as all gene pairs.

**Figure S7:**
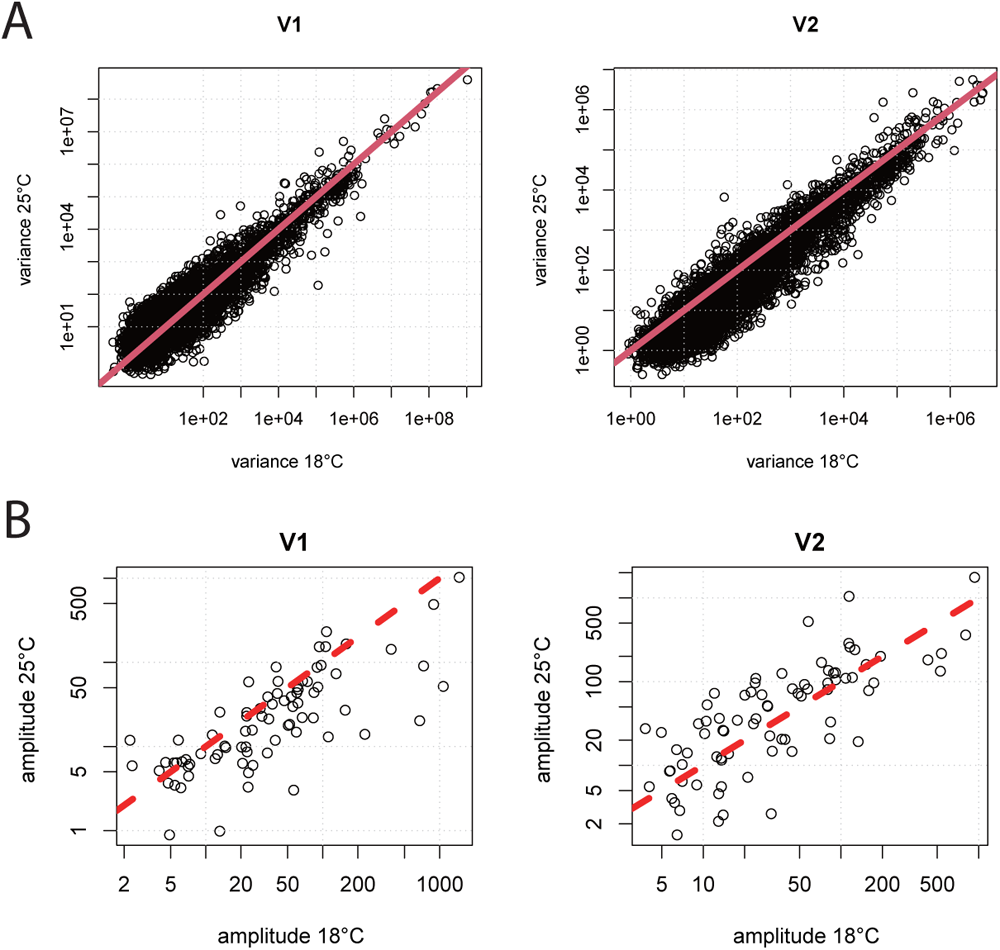
Comparison of variance and amplitude in 25*^◦^*C relative to 18*^◦^*C in the two datasets. A: Gene expression variance for all genes passing filtration in V1 and V2. B: Oscillation amplitude of genes cycling under both temperatures in V1 and V2. In all plots, red lines indicate *y* = *x*.

